# Evidence for noisy oscillations of cAMP under nutritional stress condition in budding yeast

**DOI:** 10.1101/2023.01.23.524687

**Authors:** Sonia Colombo, Maddalena Collini, Laura D’alfonso, Giuseppe Chirico, Enzo Martegani

## Abstract

The Ras/cAMP/PKA pathway is one of the best studied signalling pathway in the budding yeast that regulates cellular responses to nutrients availability and stress. The cAMP levels and the PKA activity are also subjected to a strong negative feedback that operates either through the activity of the phosphodiesterase Pde1 and also on the amount of Ras2-GTP. We have previously made and simulated a dynamic model of the whole pathway and our results suggest the existence of stable oscillatory states that depend on the activity of the RasGEF (Cdc25) and RasGAP (Ira proteins) (Pescini et al. Biotechnol Adv 30, 99-107, 2012). Stochastic oscillations related to activity of the pathway were reported by looking at the nuclear localization of the trascription factors Msn2 and Msn4 (Gamedia-Torres et al. Curr Biol 17, 1044-9, 2007). In particular Medvedik et al. (PloS Biol 5, 2330-41, 2007) reported stable oscillations of the nuclear accumulation of Msn2 in condition of limited glucose availabiliy.

We were able to reproduce the periodic accumulation of Msn2-GFP protein in yeast cells under condition of limiting glucose, and we tried to detect also in the same condition oscillations of cAMP levels in single yeast cells. We used a specific Fluorescence Resonance Energy Transfer (FRET) sensor based on a fusion protein between CFP-EPAC-YFP expressed in yeast cells. The FRET between CFP and YFP is controlled by cAMP concentration. This sensor allows us to monitor changes in cAMP concentrations in single yeast cell for a relative long time and a peak of cAMP was normally detected after addition of glucose to derepressed cells (Colombo et al. Biochem Biophys Res Commun 487, 594-99, 2017). Using this method we were able to detect noisy oscillations of cAMP levels in single yeast cells under condition of nutritional stress caused by limiting glucose availability (0.1%). We used Spectral analysis to discriminate between true oscillations and random noise. The oscillations were characterized by period of about 4-5 min, close to that observed for Msn2-GFP oscillations.

## Introduction

In the yeast *Saccharomyces cerevisiae*, the cAMP/PKA pathway plays an important role in the control of metabolism, stress resistance, proliferation (1,2,3,4,5). The central component of this pathway is the adenylate cyclase whose activity is controlled by two G-protein systems, the Ras proteins and the G α protein Gpa2 (6, 7). Cyclic AMP is synthesized by adenylate cyclase, encoded by *CYR1* gene, and induces the activation of the cAMP-dependent protein kinase A (PKA). In turn, PKA phosphorylates a variety of proteins involved in key cellular processes. The whole signaling cascade is tightly regulated and experimental evidences indicate that multiple feedback mechanisms operate within the pathway by the generation of a complex interplay between the cascade components (8, 9,10).

Ras proteins are positively controlled by the activity of Cdc25, that stimulates the GDP-GTP exchange, and negatively regulated by Ira1 and Ira2, that stimulate the GTPase activity of Ras proteins. The degradation of cAMP is governed by phosphodiesterases that constitute the major feedback mechanism in the pathway (8), although Colombo *et al*. (9) demonstrated that the feedback inhibition mechanism acts also by changing the Ras2 protein activation state. PKA can phosphorylate Cdc25, reducing its exchange activity (11), and Ira protein likely regulating the Ras-GAP activity (12, 13).

We have previously developed and simulated a dynamic model of the whole pathway and our results suggest the existence of stable oscillatory states that depend on the activity of the RasGEF (Cdc25) and RasGAP (Ira proteins) (10). Moreover we found that stable oscillatory regimes of intracellular cAMP levels can be established when a feedback operating on Ira protein is activated and the insurgence of the oscillations is deeply affected by the intracellular GTP/GDP ratio linking the transition between stable steady states and oscillations to a reduced nutritional conditions. Sustained oscillations of cAMP level have never been observed in budding yeast, although in some cases damped oscillations were reported after addition of glucose to glucose-derepressed cells (14) on yeast populations.

However the existence of sustained oscillatory states can be inferred in indirect way by looking to the nuclear localization of the transcription factors Msn2 and Msn4. In fact stochastic oscillations related to activity of the PKA pathway were reported by looking at the nuclear localization of the transcription factors Msn2 and Msn4 (15). In particular Medvedik et al. (16) reported stable oscillations of the nuclear accumulation of Msn2 in condition of limited glucose availability.

Although the cAMP/PKA pathway has been extensively studied in yeast and both upstream and downstream elements are known, the changes in cAMP and in the activity of this pathway were usually measured in cell populations for a very short time and data on the spatiotemporal variation of cAMP and PKA activity in single cells are till now lacking. Some years ago Nikolaev *et al*. developed Fluorescence Resonance Energy Transfer (FRET) probes Epac-based for monitoring cAMP levels *in vivo* in single mammalian cells (17). These sensors consist of part of the cAMP-binding protein Epac1 or Epac2 sandwiched between cyan and yellow fluorescent proteins (CFP and YFP). The construct unfolds upon binding of the second messenger cAMP to the Epac moiety and cAMP increases are thus easily followed as a drop in FRET (17). We previously adapted these Epac-based probes for expression in yeast vectors and we found a specific FRET signal clearly related to changes of cAMP on single yeast cells using a confocal microscope (18). Here we used this FRET based cAMP sensor to evidence the presence of sustained oscillation of cAMP in single yeast cells in condition of limiting glucose (0,1%), i.e under a nutritional stress.

## Materials and Methods

### Yeast strains and media

Strains used in this study: SP1 (*MATa his3 leu2 ura3 trp1 ade8 can1*) [19]; BY4741-YMR037C (*MATa his3 leu2 ura3 MSN2*-GFP) (Invitrogen). Yeast cells were grown in synthetic complete media (SD) containing 2% glucose, 6.7 g/l YNB w/o aminoacids (supplied by ForMedium™, United Kingdom) and the proper selective drop-out CSM (Complete Synthetic Medium, supplied by ForMedium™, United Kingdom) at 30°C in shaken flasks. Culture density was measured with a Coulter Counter (Coulter mod. Z2) on mildly sonicated samples.

### Plasmids

To obtain the pYX212-YFP-EPAC2-CFP and the pYX242-YFP-EPAC2-CFP vectors we used the following strategy. The YFP-EPAC2-CFP fragment, obtained digesting the pcDNA3-YFP-EPAC2-CFP construct (kindly provided by Dr. V.O. Nikolaev, University of Wuerzburg, Germany) (17) with *Xho*I and *Hind*III, was ligated into the expression vectors, pYX212 and pYX242, digested with the same enzymes (18).

### Fluorescence microscopy and FRET determination

Cells were grown in medium containing 2% glucose at 30 °C till exponential phase, collected by centrifugation, washed 2 times with SD medium containing 0.1% glucose and suspended in SD + 0.1% glucose medium at a density of 5 × 10^7^ cells/ml and incubated at 30 °C for at least 1 h. Subsequently, 40 μl of cell suspension were seeded on concanavalin A (ConA) (Sigma-Aldrich, Milano, Italy)-coated cover glass for 10 min (18,20). The cover glass was washed four times using 1 ml of SD+ 0,1% glucose medium, mounted on a custom chamber and covered by 500 μl of the same medium. Time stacks of images (512×512 pixels, typical field of view 150 μm x 150 μm, 400 Hz scanning frequency) were acquired by means of a Leica SP5 confocal microscope (Leica BLA Germany) through a 40X oil objective (HCX PL APO CS 1.30). The pinhole was set at 150-170 μm in order to detect a higher signal from each cell and to avoid losing the focal plane in long time acquisitions. CFP was excited at 458 nm, its emission detected in the range 468-494 nm and YFP emission was detected in the range 530-600 nm. The acquisition (1 image every 2.6 sec) started without interruption for at least 60 minutes. We have previously shown that under these conditions the fluorescence signal was stable for a long time with a minimal photobleaching (18) For each sample, image time series have been acquired selecting field of view populated with more than 50 cells. After acquisition, data were analyzed by means of the Leica Application Suite Software (Leica Microsystem, Germany). A ROI (Region of Interest) was selected including each cell and the CFP and YFP fluorescence signals in the two acquisition channels have been saved together with their ratio versus time. In this way, both single cell behavior and average values were calculated for each sample. The raw data were then further elaborated with Excel ™.

For the analysis of Msn2-GFP nuclear localization yeast cells (BY4741-YMR037C), expressing the Msn2-GFP fusion protein were grown till exponential phase in synthetic medium (SD + 2% glucose). Cells were collected by centrifugation (5 min at 3000 rpm), washed 2 times with SD medium containing 0.1% glucose and suspended in SD + 0.1% glucose medium at a density of 5 × 10^7^ cells/ml. After incubation of 60 min at 30°C aliquot (20 μL) of cell suspension was seeded on a coverslip coated with concanavalin-A (18,20) and put on top of a Thoma chamber. Images were acquired with a Nikon Eclipse 90i fluorescence microscope equipped with a 60× oil immersion objective with GFP adapted filters and photographs were taken at specific time intervals. To avoid bleaching, the fluorescence images were acquired for 1 s every 2 minutes and the shutter was kept off in the meantime. Images were analyzed with ImageJ software (https://imagej.nih.gov) in order to calculate the ratios of average nuclear intensity versus average cytoplasmic intensity of green fluorescence.

### Analysis of periodicity

Spectral analysis of raw data was done using the software Kyplot 5 ™, running on a PC (Windows 7) (http://www.kyenslab.com/en/kyplot.html). A noise reduction (smoothing) was sometimes done using a moving average of 20 experimental points in Excel. The raw or smoothed data were also used for Recurrence analysis (21) (Visual Recurrence Analysis Software: http://www.visualization-2002.org/VRA_MAIN_PAGE_.html). Simulated cAMP data were obtained with the software BioSimWare (10), using a personal computer running Windows XP. All stochastic simulations were performed by exploiting the tau-leaping algorithm (22).

## Results and discussion

### Nutritional stress condition induces stochastic periodic nuclear localization of Msn2 protein

We have previously developed and simulated a dynamic model of the Ras/cAMP/PKA pathway in budding yeast and our results suggest the existence of stable oscillatory states that depend on the activity of the RasGEF (Cdc25) and RasGAP (Ira proteins) (10). An example of these oscillatory regimes is shown in Supplementary material (Fig. S1).

In addition we have shown that cAMP levels and PKA activity can be measured in single yeast cells using specific FRET probes (18), therefore we investigated the possibility to evidence cAMP oscillations in single yeast cells using the CFP-EPAC2-YFP probe (17,18). In preliminary experiments we tested the conditions required to induces oscillations in the nuclear localization of Msn2 protein. To do this we used an yeast strain that expresses a fusion of Msn2 with eGFP, in this strain the green fluorescence was diffused in the cytoplasm when the cells were growing in glucose medium and relocalized in the nucleus after glucose starvation (16). Under condition of limiting glucose availability (0,1% glucose, i.e. in a nutritional stress) we were able to evidence the insurgence of asynchronous stochastic oscillations (Fig.1, and Supplementary movie 1) in agreement with the data reported by Medvedik et al. (16). As shown in Fig.1 each cell behaves in a different and asynchronous way with a periodicity in the range of 5-10 min.

**Fig. 1.**
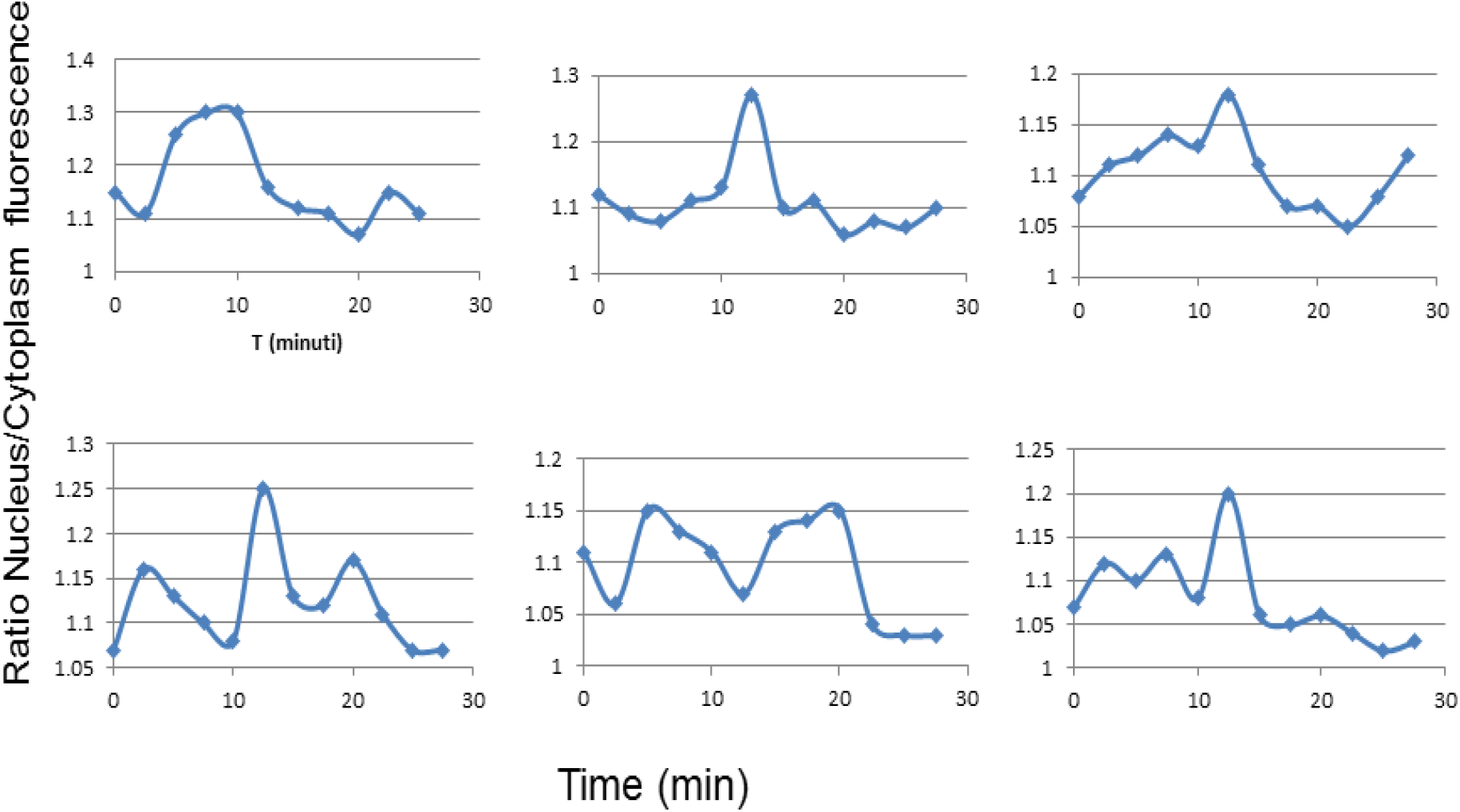
Time-course of localization of Msn2-Gfp protein in the nucleus of six yeast cells.

### Nutritional stress induces noisy oscillation of cAMP in single yeast cells

To evaluate the presence of oscillations of cAMP levels in yeast cells we used a strain that expresses the FRET probe CFP-Epac2-YFP (18). This strain was grown exponentially in 2% glucose synthetic medium (SD) and then the yeast cells were resuspended in the same medium containing a reduced amount of glucose (0.1% w/v). After 1 hour the cells were seeded on a cover glass precoated with ConA and incubated in a small chamber with the same low glucose medium as reported previously (18). The chamber was then mounted on a Confocal Inverted Microscope and time stacked images were acquired as reported in Materials and Methods. An example of the acquired images is shown in Supplementary Fig. S2. After acquisition of a time series of images a Circular Region (ROI) was selected including each cell and the CFP and YFP fluorescence signals in the two acquisition channels was measured and saved together with their ratio versus time. In this way, both single cell behavior and average values were calculated for each sample.

We collected time series for more than 100 single cells, and we found a lot of random noise in the fluorescence ratio used for FRET analysis (YFP/CFP fluorescence); several raw data are shown in the Fig 2 We used mainly two different approaches to filter the noise in order to evidence the eventual presence of oscillations, a simple moving average window that is enough to reduce the noise and a more sophisticated Spectral Analysis of raw data that evidences the presence of low frequency components well separated from the higher frequency noise. Both methods evidenced an high heterogeneity in the behavior of single yeast cells: in many of them only a noise was apparent without any specific low frequency oscillations (examples are shown in Fig. 3), in another group of single cells low frequency oscillations with a period in the range of 4-6 minutes were detectable immediately after the start of data acquisition (Fig. 4), finally in some other cells a more interesting behavior was observed, after a variable period without oscillations, at a given point low frequency oscillations appeared (Fig. 5 and 6). In all these cases the FRET signal steadily increased during the first non-oscillatory period suggesting that the intracellular level of cAMP decreased till the insurgence of oscillations. In these cases the spectral analysis performed on the whole raw data failed to evidence the presence of low frequency oscillation, that however were well evident if the analysis is performed on the second half of the data (Fig. 5 and 6).

**Fig. 2.**
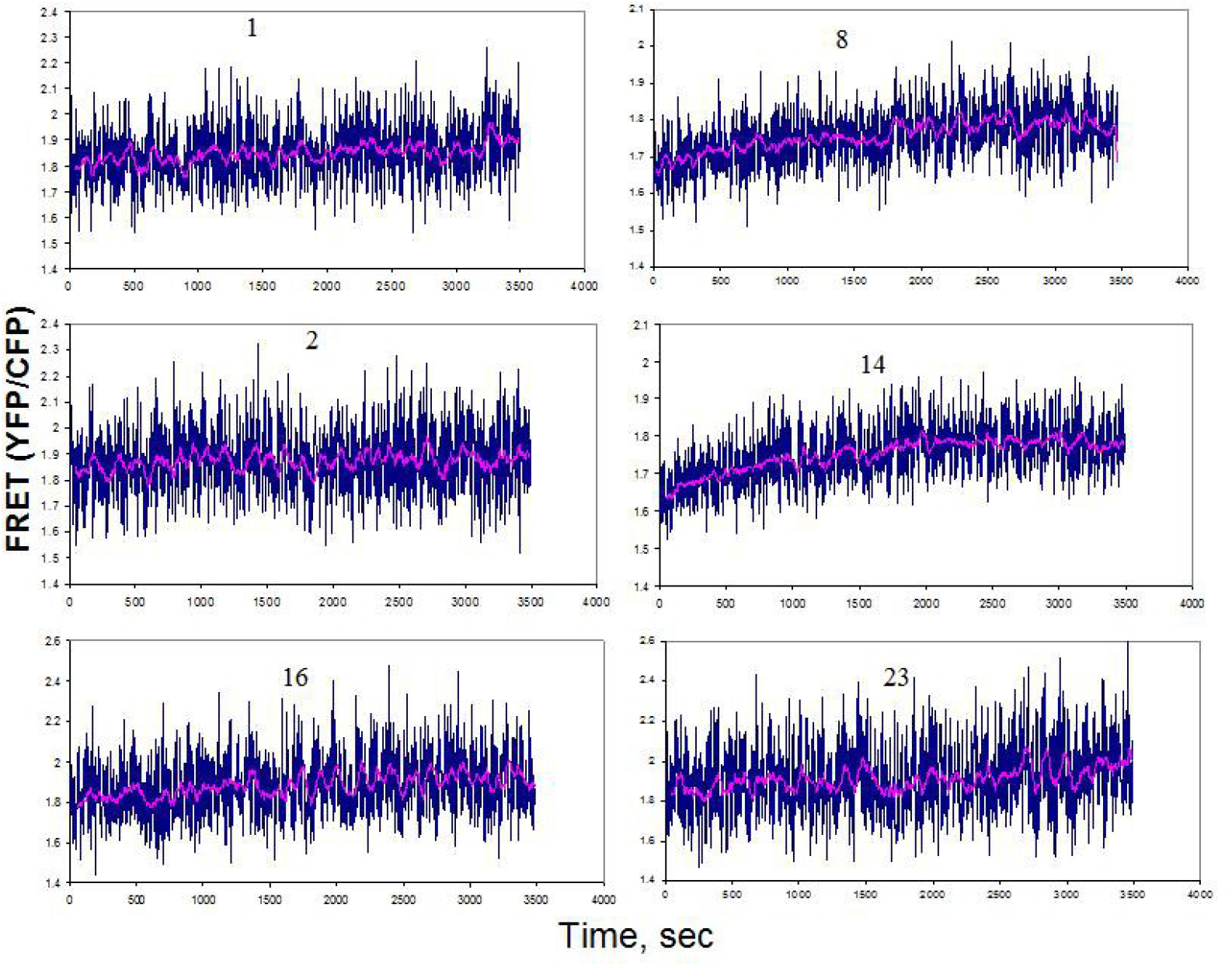
Example of FRET measured in single yeast cell every 2.6 seconds for a total of for 3490 sec. The raw data are reported in blue, while a moving average of 20 points is reported in pink. The number (1,2,8 etc..) indicated the ROI of the measured cell (see Supp Fig.2)

**Fig. 3.**
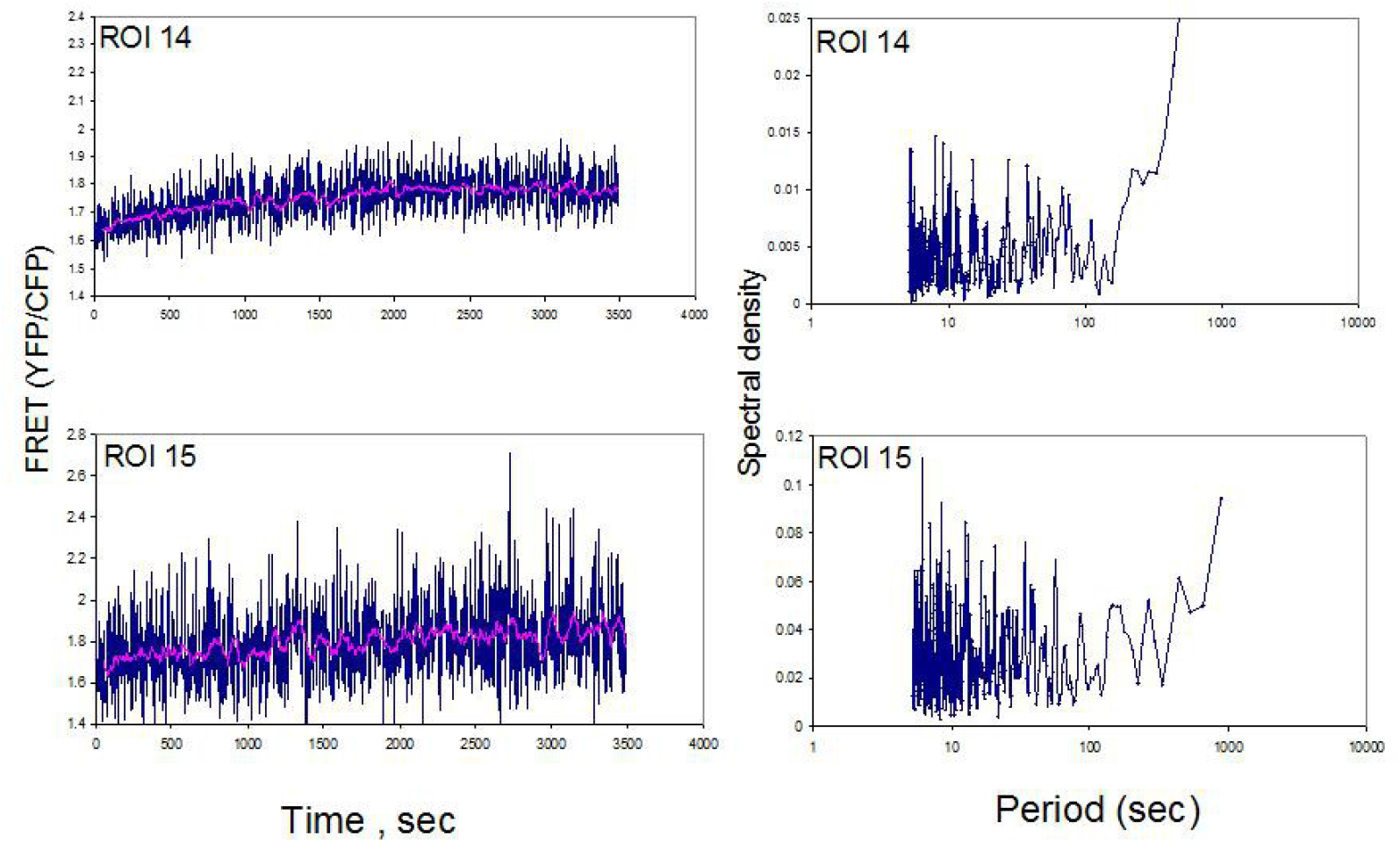
FRET time course of two cells that fail to exihibit significant low frequency oscillations. The spectral analysis does not evidence any low frequency components emerging over the noise.

**Fig. 4.**
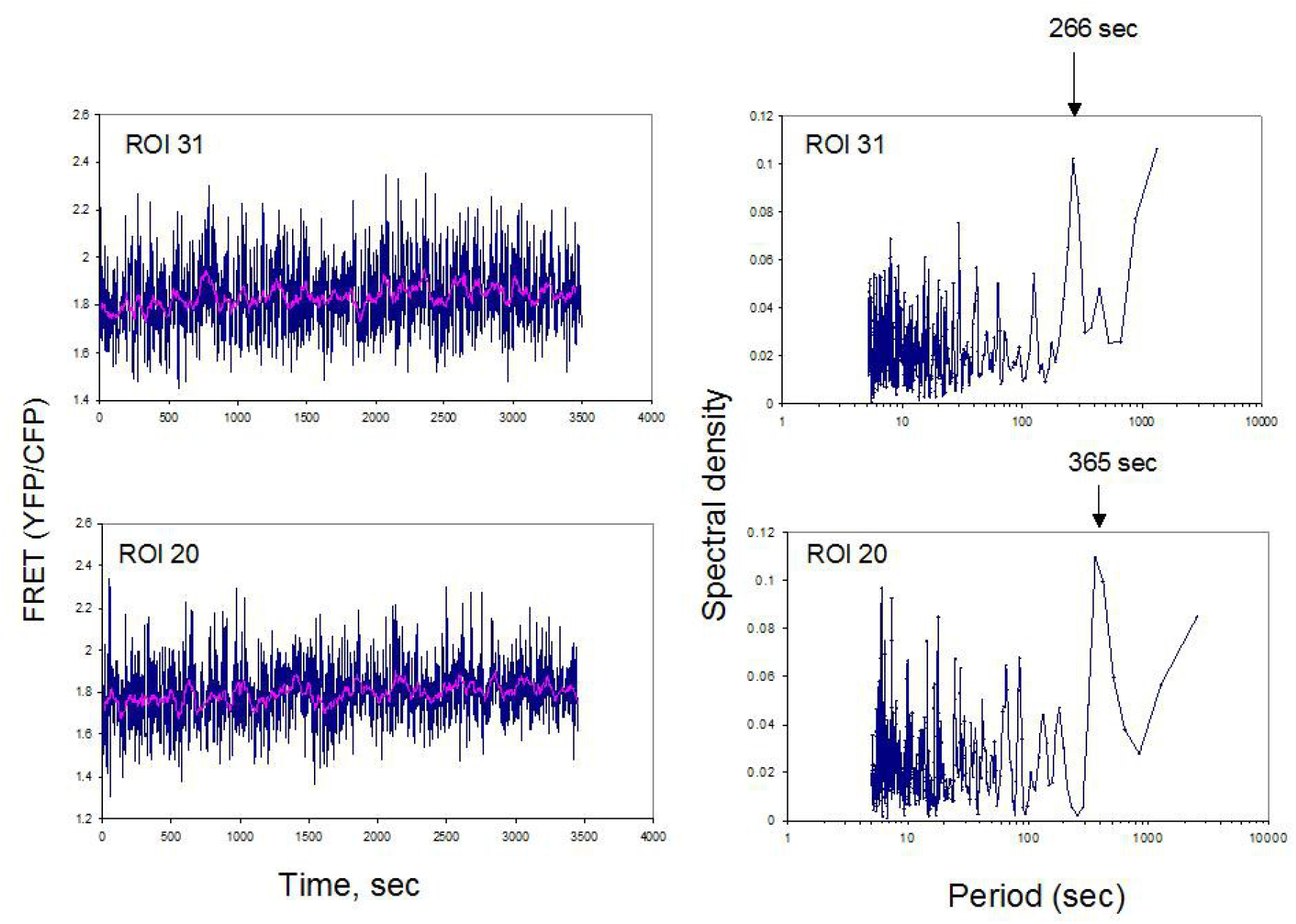
Example of cells that exhibits a noise oscillation from the beginning. The spectral analysis evidenced a low frequency component with a period of in the range of 4-6 min.

**Fig. 5.**
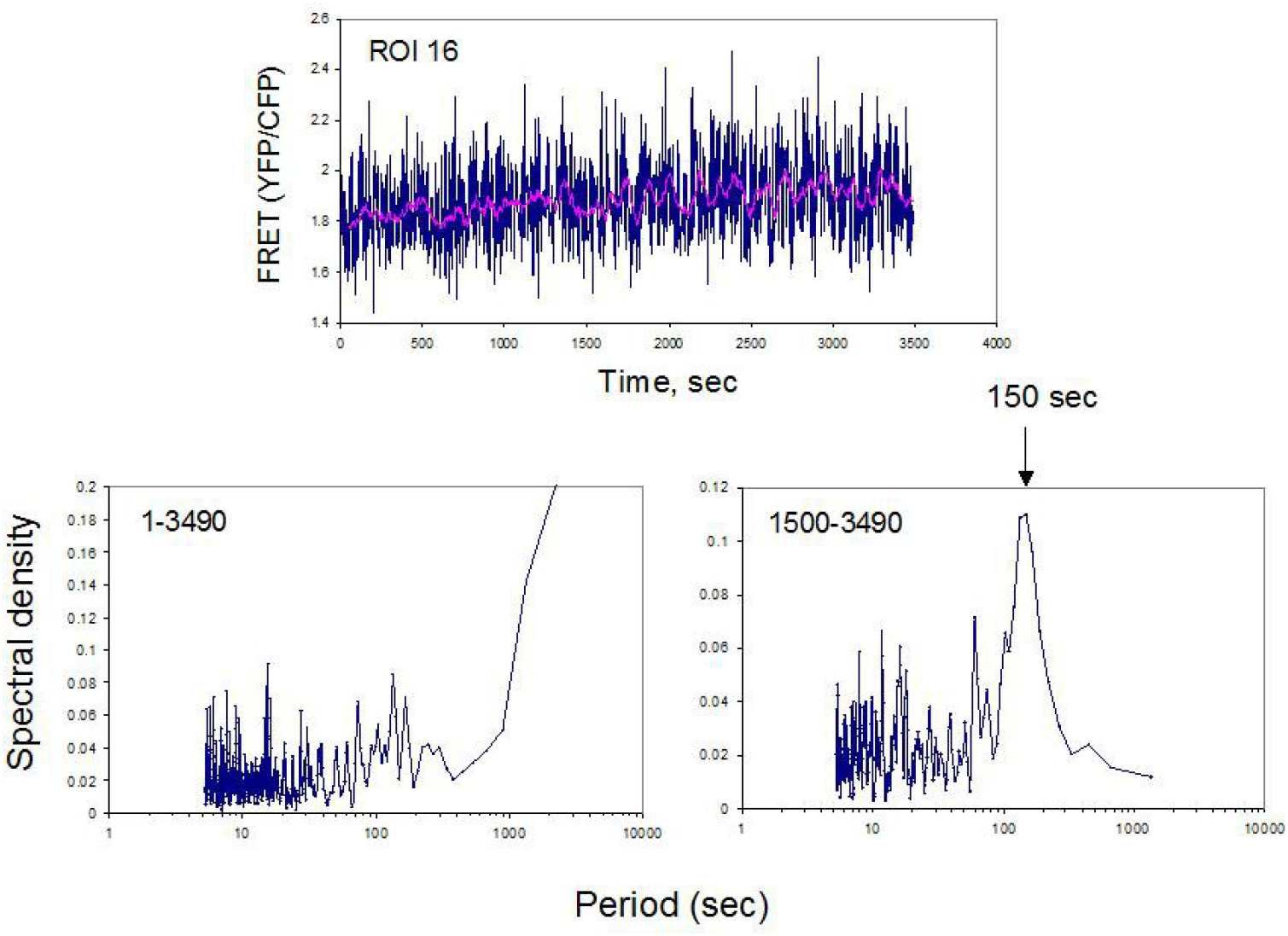
Example of a cell that exhibits noise oscillations only during the second part of the experiment (after 25 minutes)

**Fig. 6.**
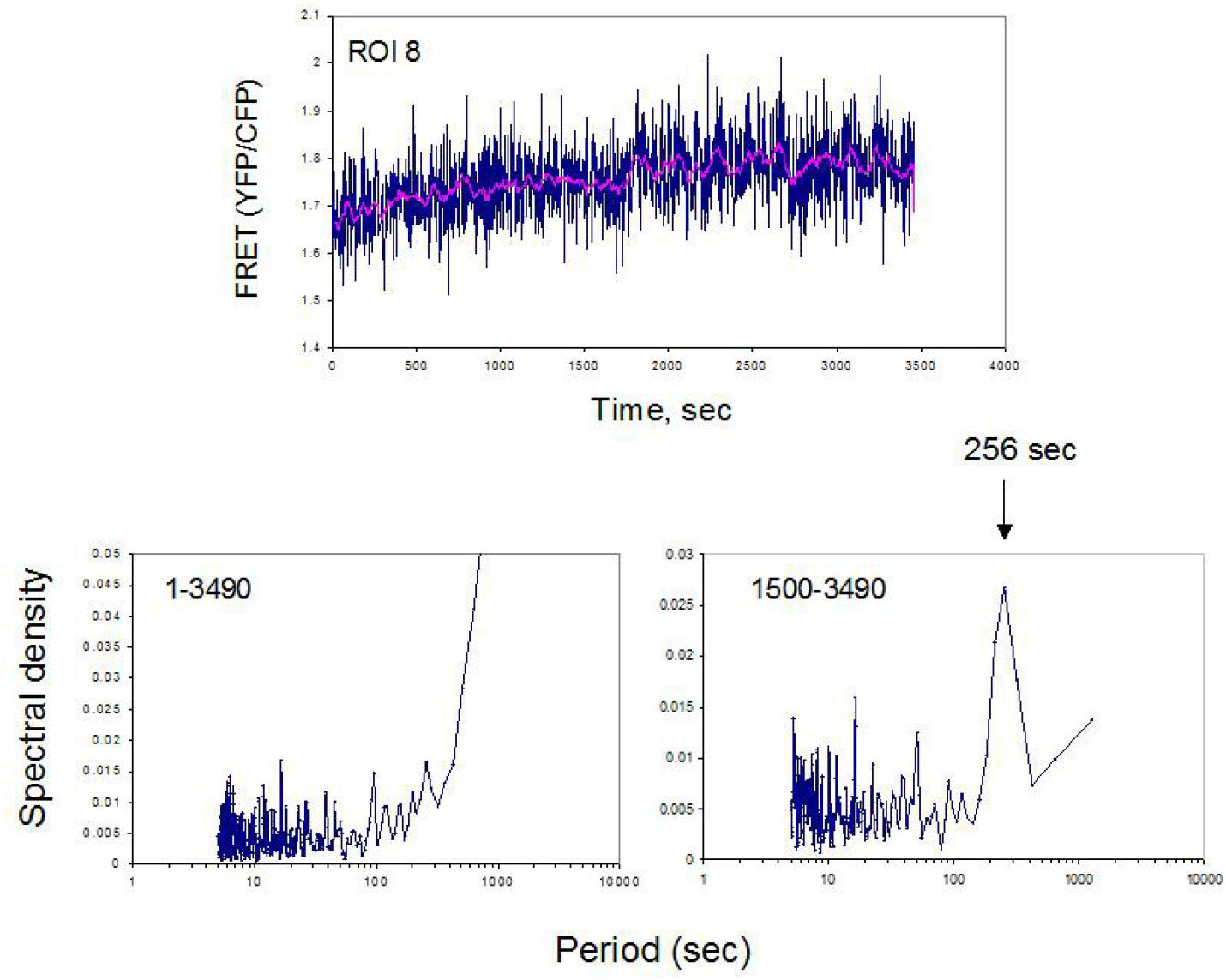
The cell identifyed as ROI 8 showed clear noise oscillations starting after 26 min. The period is of about 4 min

An example of this behavior is also well evident for the cell analyzed as Roi8 (Fig.6). The presence of oscillations in the second half of the data (points from 1500 to 3490 sec) is barely evident on raw data, but it is clear after a smoothing (using a moving average window of 20 experimental points). The apparent period is of 4 min, in agreement with spectral analysis performed on the same raw data. The reverse of FRET (i.e CFP/ YFP fluorescence signal) is directly related to intracellular cAMP concentration, that steadily decreases in the first 600 points (equivalent to 26 min) and then starts to oscillate in the second part for at least 30 min. (Supplementary Fig. 3)

Taking in account the affinity of EPAC2-Camp for cAMP (Kd= 0.8-1 μm) (17, 23) we estimated that intracellular cAMP concentration oscillates between 0.2 and 0.8 μM (about 6000-25000 molecule/cell) in the range reported in ref. (14) (Supplementary Fig. 4).

A similar behavior was observed for other cells (Roi 16 for example) and the insurgence of an oscillatory state can be also evidenced by a Recurrence Analysis software as shown in fig. 7 where it is well visible a change of pattern and the appearance of many lateral diagonals indicating a periodicity.

**Fig. 7.**
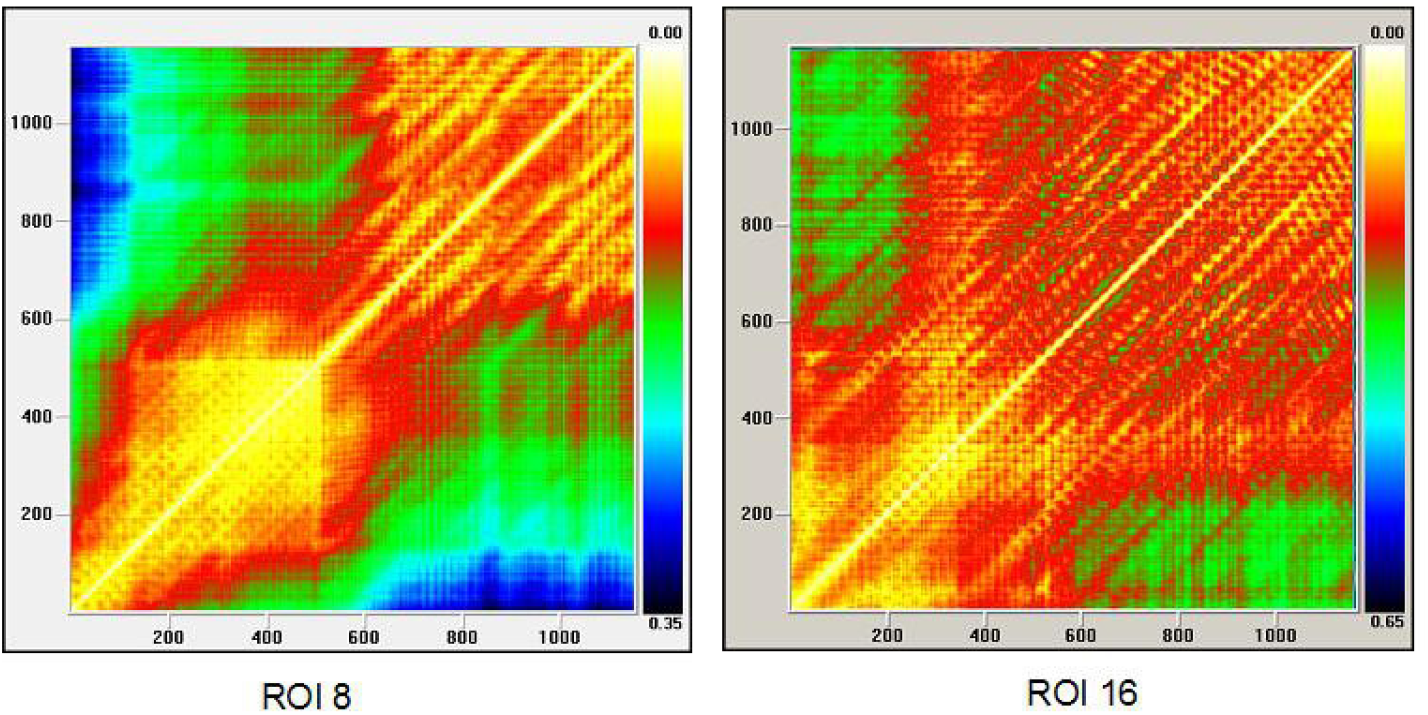
Recurrence analysis performed with the software -Visual Recurrence - VRA (: http://www.visualization-2002.org/VRA_MAIN_PAGE_.html) using the smoothed data (average 20) of the cells ROI 8 and ROI 16. It is well evident a change in the pattern in the second part of the data in correspondence with the insurgence of the oscillations

## Conclusions

This is the first time that spontaneous oscillations in cAMP levels were observed in budding yeast. The oscillations were simultaneosly observed in several yeast cells under conditions of nutritional stress (i.e. low glucose availability). Each cell behaves in a different way, i.e. many cells did not present any oscillations, others presented oscillations from the beginning of the experiment, while many others presented oscillations during the course of the experiment. About 30% of the observed cells showed an oscillatory pattern, however the period of the oscillations varied from 3 to 6 minutes and the oscillation were not sinchronous and in any case the oscillations were partially masked by an high frequency noise (white noise??). This oscillatory behaviour of the cAMP/PKA pathway was predicted by simulations performed with a stochastic model of this pathway (10) and could explain the nucleo-cytoplasmic shuttle of Msn2 and Msn4 transcription factors (15-16).

## Acknowledgements

We thank Dr. V.O. Nikolaev, University of Wuerzburg, Germany for providing the pcDNA3-YFP-EPAC2-CFP construct. This work was supported by FAR-University of Milano-Bicocca grants to E.M. and S.C., by Program Sys-BioNet, Italian Roadmap Research Infrastructure 2012 grant to S.C. and by fundings from University of Milano-Bicocca to M.C. and G.C. (“large infrastructures 2011 and “Unimib competitive grant”)

## Supplementary data

**Supplementary Fig. S1:**
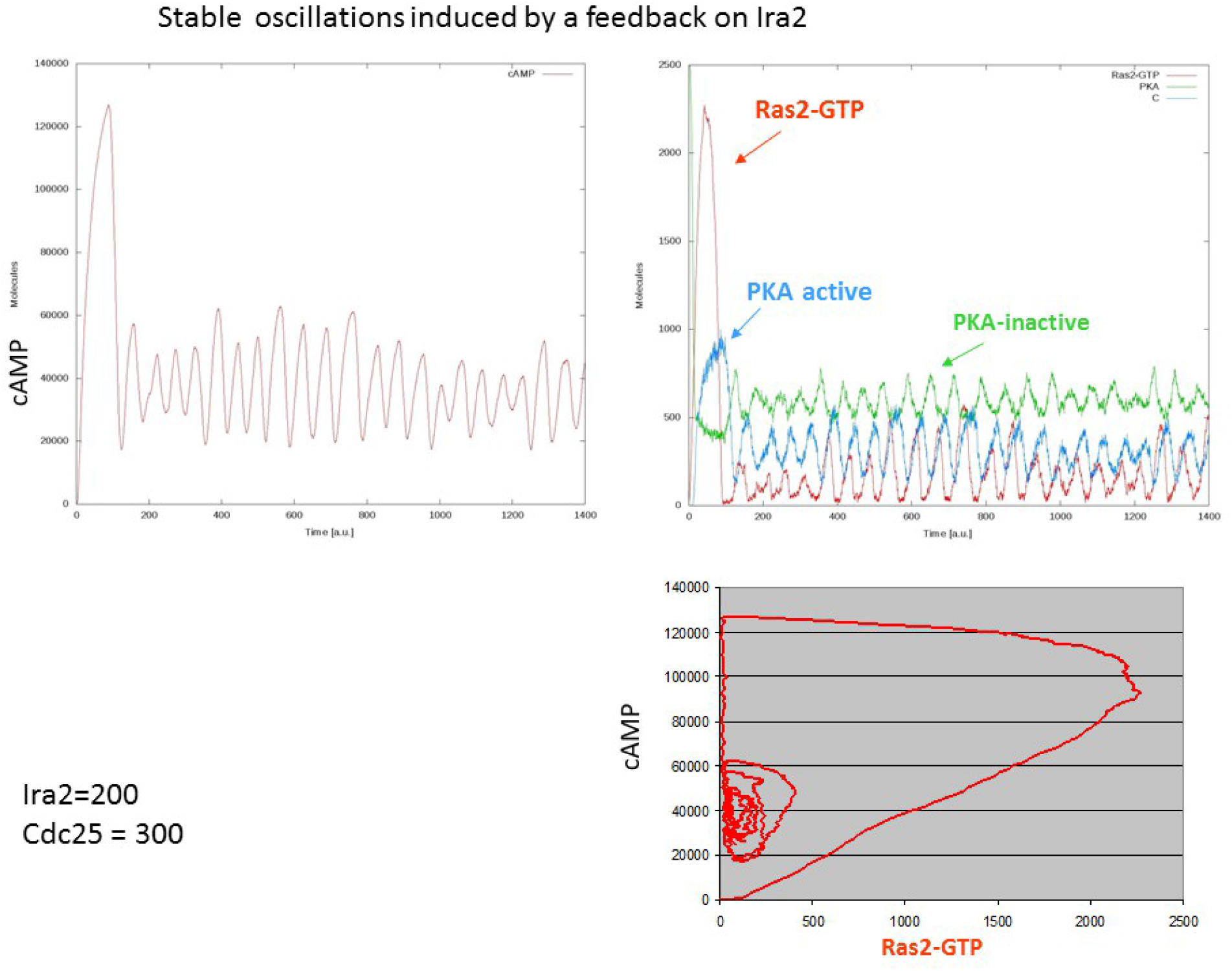
Simulation of cAMP molecules in single yeast cell. The simulation was done with the software developed in ref 10 and 22 assuming 200 molecules/cell of Ira2 and 300 of Cdc25

**Supplementary Fig. S2:**
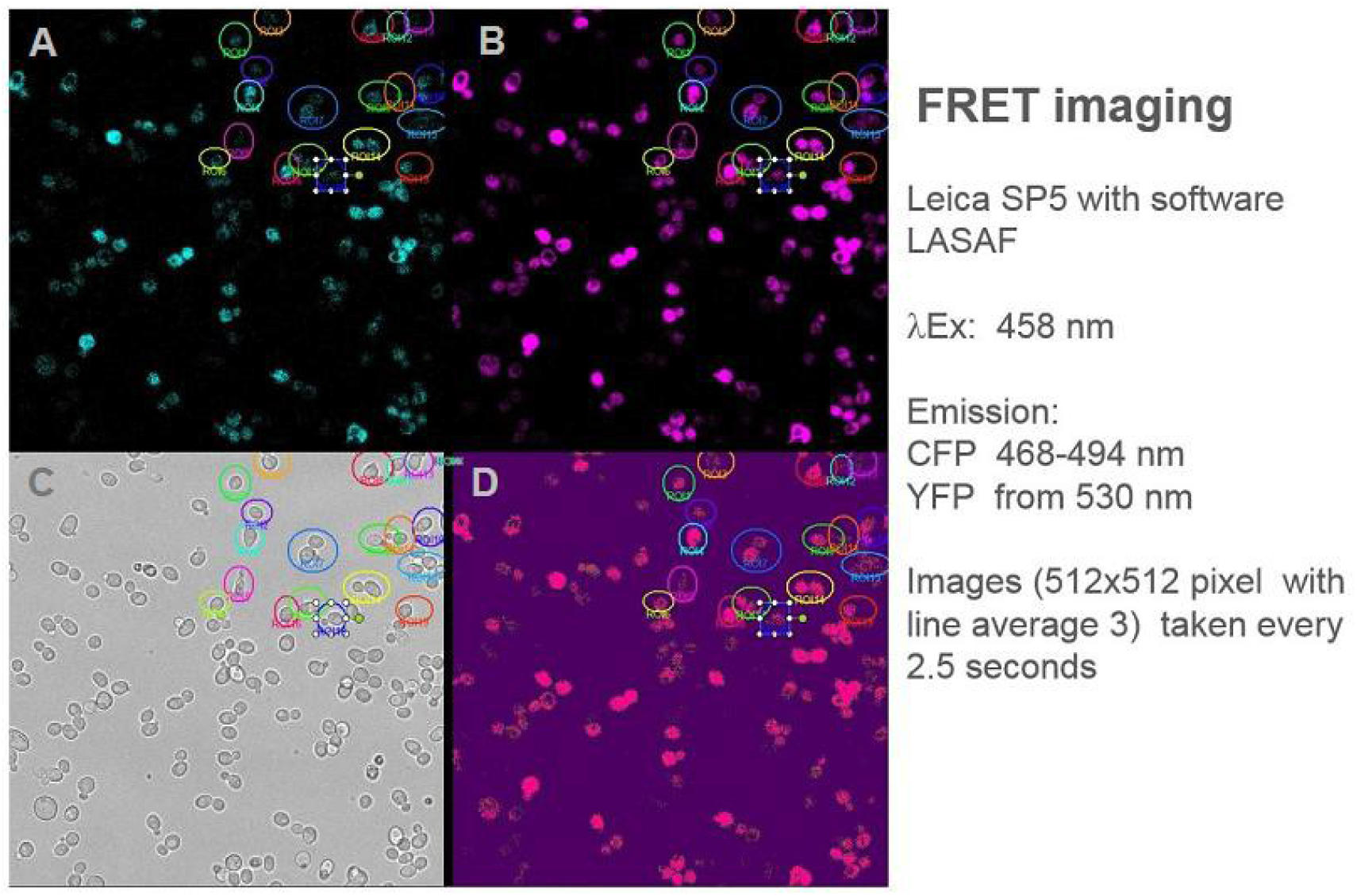
Example of imaging of yeast cells expressing the cAMP-probe: A) Fluorescence of CFP; B) Fluorescence of YFP; C) Transmission light image; D) Ratio YFP/CFP; the circle are the ROI selected

**Supplementary Figure S3:**
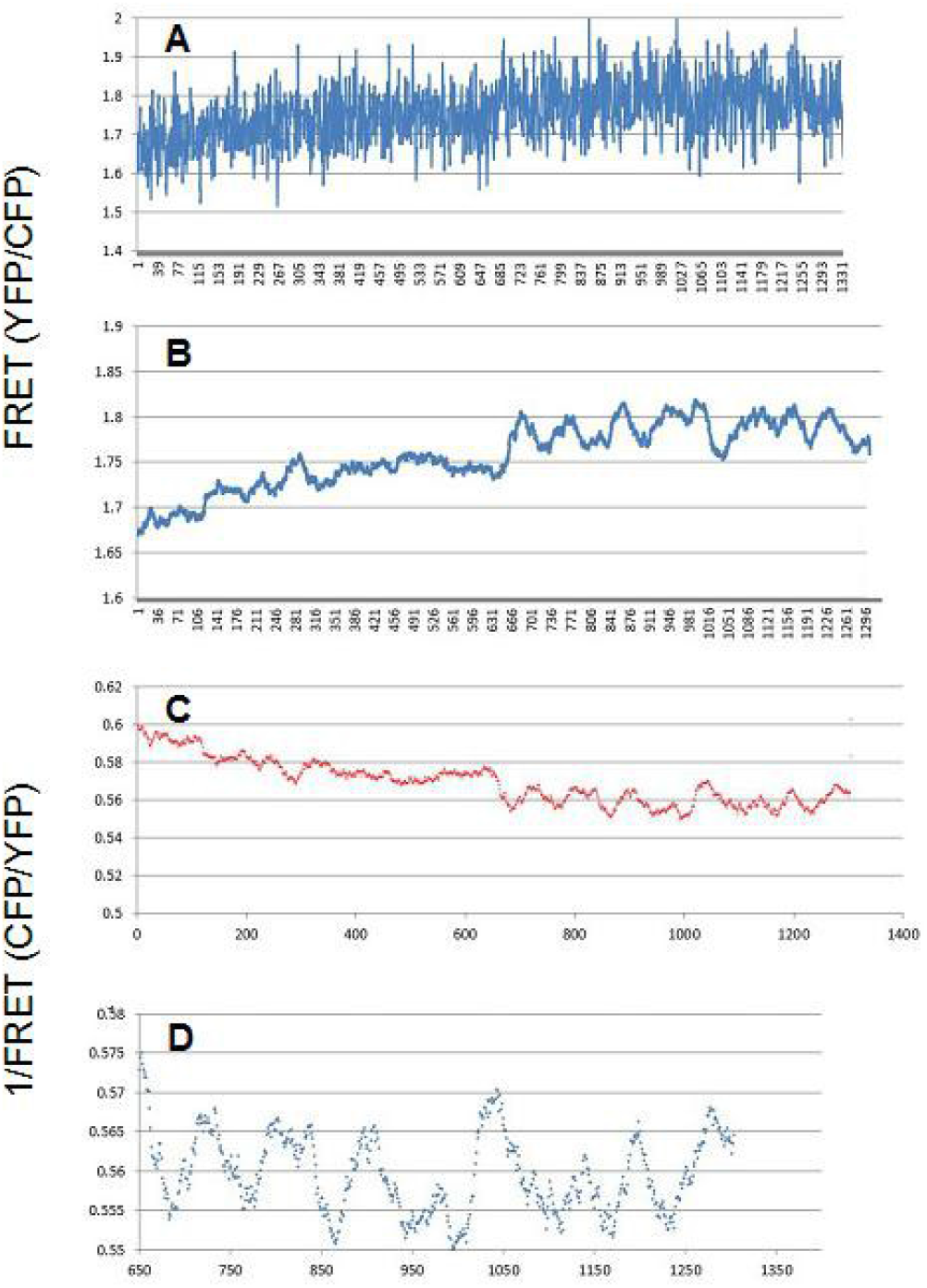
The data are related to the cell named ROI8: A) raw YFC/CFP ratio; B) denoising with an moving average of 20 experimental points; C) Moving average of the reverse of FRET (CFP/YFP) ratio that is related to the intracellular level of cAMP; D) enlargment of the second part of the data reported in (C) in order to better evidence the oscillations that started after 26 minutes.

**Supplementary Figure S4:**
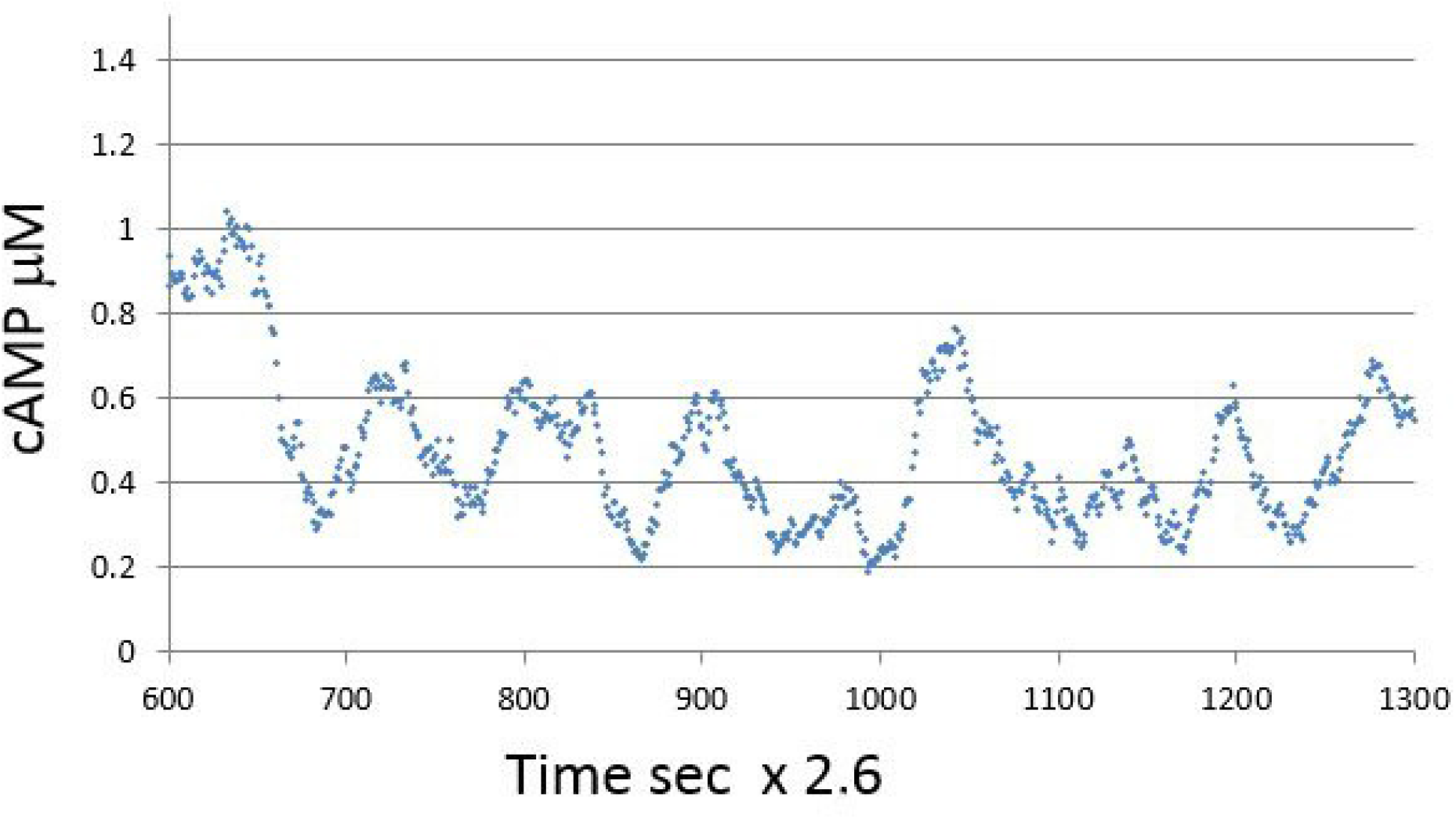
Intracellular concentration of cAMP extimated from FRET data of Supplementary Fig. S3. We used the formula reported in ref (23), assuming an Hill coefficient of 1 and a Kd of 1 μM

